# SAI: A Python Package for Statistics for Adaptive Introgression

**DOI:** 10.1101/2025.04.19.649497

**Authors:** Xin Huang, Simon Chen, Josef Hackl, Martin Kuhlwilm

## Abstract

Adaptive introgression is an important evolutionary process, yet widely used summary statistics—such as the number of uniquely shared sites and the quantile of the derived allele frequencies in such sites—lack accessible implementations, limiting reproducibility and methodological clarity. Here, we present SAI, a Python package for computing these statistics, and apply it to three datasets. First, using the 1000 Genomes Project data, we replicated previously reported candidate regions and identified additional ones, including a region detected by studies using supervised deep learning. Second, reanalysis of a Lithuanian genome dataset revealed no candidates in the HLA region. Finally, we investigated bonobo introgression into central chimpanzees and identified a candidate region that overlaps a high-frequency Denisovan-introgressed haplotype block reported in modern Papuans—an intriguing co-occurrence across divergent lineages. Discrepancies with prior results highlight the importance of transparent and reproducible analysis workflows, especially as machine learning becomes increasingly prevalent in evolutionary genomics.

---

Adaptive introgression refers to the transfer of genetic material between distantly related lineages through gene flow, increasing the fitness of the recipient lineage. To detect candidate genomic regions of adaptive introgression, some studies primarily utilized *ad-hoc* approaches—first applying methods to detect natural selection and introgression separately, then identifying their intersection as candidate regions (Fijarczyk and Babik 2015; Nye et al. 2018; Zhang X et al. 2023). Furthermore, supervised machine learning methods, such as genomatnn, MaLAdapt, and ERICA, have been introduced specifically for detecting loci of adaptive introgression (Gower et al. 2021; Zhang X et al. 2023; Zhang Y et al. 2023). Meanwhile, dedicated summary statistics for this purpose have been proposed by Racimo et al. (2017) (see Supplementary Table S1), such as the number of uniquely shared sites (*U* statistic) and the quantile of the derived allele frequencies in such sites (*Q* statistic). These statistics remain invaluable, particularly in cases where the demographic model is unknown (Gower et al. 2021; Romieu et al. 2024). Despite their utility, these statistics lack a readily available implementation, requiring researchers to develop custom scripts for their analyses, which could harm accessibility and reproducibility of research. To address these gaps, we developed SAI (version 1.0.1), a Python package designed for calculating the *U* and *Q* statistics, providing a robust framework for future studies on adaptive introgression (Supplementary Material). We excluded the *R*_*D*_ statistic proposed by Racimo et al. (2017), as its implementation may require excluding zero-denominator cases and introduce systematic bias (Supplementary Material).

We first applied SAI to the 1000 Genomes Project Phase 3 dataset (The 1000 Genomes Project Consortium 2015), as previously reported in Racimo et al. (2017) (see Supplementary Material). Here, the Altai Neanderthal and Denisovan genomes were used as source or donor populations, that is, the populations presumed to have contributed introgressed material (Huang et al. 2022). Candidate regions were identified based on the overlap of outliers from both the *U*50 and *Q*95 statistics, computed using derived allele frequencies (Supplementary Table S2). These regions were then annotated using ANNOVAR (version 2022Oct05), with annotations from RefSeq, dbNSFP v4.2c, and dbSNP build 150 (Sherry et al. 2001; Wang et al. 2010; O’Leary et al. 2016; Liu et al. 2020). We successfully replicated several previously reported candidate regions, including those in *BNC2*, a gene associated with human pigmentation and may subject to natural selection in modern Europeans (Cheng et al. 2021; Huang et al. 2021). We also identified additional candidate regions, including one on chromosome 20 that was not reported by Racimo et al. (2017) but was later detected using supervised deep learning (Gower et al. 2021). However, we did not recover the region chr17:18880001–18920000 (hg19 coordinates), which Racimo et al. (2017) identified as a candidate region of Denisovan-specific introgression.

We observed discrepancies in *U*50 and *Q*95 values between our results and those reported by Racimo et al. (2017) within the same genomic regions (Supplementary Table S2). To validate our SAI results, we compared allele counts computed by our implementation with those obtained using BCFtools (version 1.21; Danecek et al. 2021) and found them to be consistent (Supplementary Material). We also noted that the original definition of the *U* statistic is not specifically based on the derived allele (Supplementary Table S1). Therefore, we extended our implementation to compute these statistics using either allele 0 or allele 1, depending on which allele meets the required conditions (Supplementary Material). Despite this adjustment, our results (Supplementary Table S3) still did not align with those of Racimo et al. (2017). We further filtered out low-quality variants in the archaic hominin genomes (Supplementary Material), but the discrepancies persisted (Supplementary Tables S4 and S5). Although the original definition of the *Q* statistic is based on the derived allele, Racimo et al. (2017) did not provide explicit details on how ancestral states were inferred when analyzing real data. A similar issue arises in MaLAdapt (Zhang X et al. 2023), which uses the *U* and *Q* statistics as input features for its machine learning algorithm (Extra-Trees classifiers). While MaLAdapt explicitly defines these statistics as based on the derived allele, the source of ancestral allele information is not specified. Since Racimo et al. (2017) did not make their code or detailed data processing steps publicly available, the precise causes of these discrepancies remain unresolved.

We then applied SAI to the Lithuanian genomes published by Urnikyte et al. (2023) to screen for candidates of adaptive introgression from Neanderthals (Supplementary Material). As in our previous study (Hackl and Huang 2025), we used BEAGLE 5.4 (version 06Aug24.a91) to impute missing genotypes, using all European populations from the 1000 Genomes Project as the reference panel (Browning et al. 2018; Browning et al. 2021). We did not identify any variants in the HLA region as candidates for adaptive introgression (Supplementary Tables S6 and S7), consistent with our earlier findings using MaLAdapt (Hackl and Huang 2025). Notably, the region chr20:62110001–62220000 (hg19 coordinates) was detected as a Neanderthal-introgressed candidate, in agreement with both our results above (Supplementary Tables S2–S5) and a previous study that reported this region in European populations from the 1000 Genomes Project (Gower et al. 2021). However, because our approach requires the allele to reach high frequency in the target population, we may miss candidates under weak or ongoing selection—particularly those maintained at low or intermediate frequencies.

Finally, we explored bonobo introgression into central chimpanzees using a curated great ape genome dataset (Han et al. 2025) with SAI, with western chimpanzees as non-introgressed reference population (Supplementary Material). Compared to human genome data, the great ape genomes showed higher values of the *U* statistic (Figure 1), indicating a substantial number of shared variants between central chimpanzees and bonobos—consistent with a previous study (de Manuel et al. 2016). Among our candidate regions (Supplementary Tables S8 and S9), we found that three genes—*ADGRL4, GALNTL6*, and *SORCS1*—had been previously reported as candidates for positive selection within bonobo-introgressed regions (Nye et al. 2018). Interestingly, we also identified the region chr14:58400001– 58450000 (hg38 coordinates), which contains *TOMM20L, TIMM9*, and *KIAA0586*, as a candidate region (Supplementary Table S9). A previous study reported that this region assigned to high-frequency Denisovan-introgressed haplotype blocks in modern Papuans (Jacobs et al. 2019).

**Figure 1.**
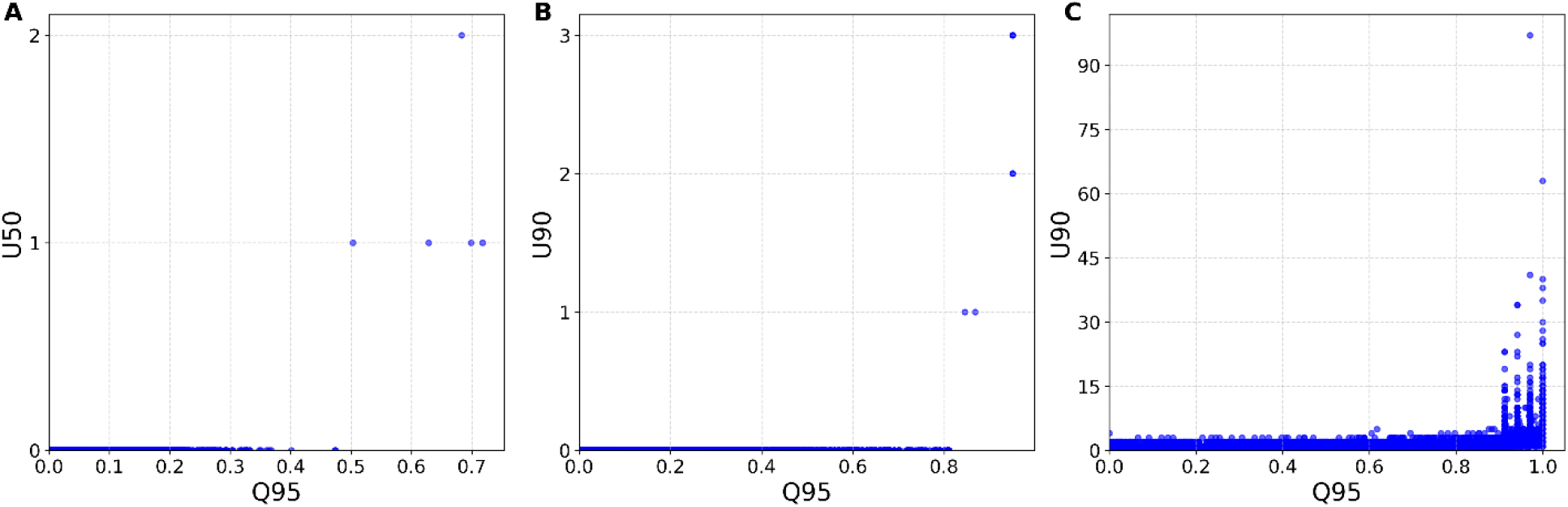
Joint distributions of genome-wide *U* and *Q* values computed using derived allele frequencies. (A) Joint distribution of *U*50 and *Q*95 values from the 1000 Genomes Project. Following Racimo et al. (2017), the filtering criteria required allele frequency < 0.01 in both the AFR and EAS populations, > 0.5 in the EUR population, and fixed in both the Neanderthal and Denisovan genomes. (B) Joint distribution of *U*90 and *Q*95 values from the Lithuanian genomes. The criteria required allele frequency < 0.01 in the YRI population, > 0.9 in the Lithuanian population, and fixed in the Neanderthal genome. (C) Joint distribution of *U*90 and *Q*95 values from genomes of the *Pan* clade. The criteria required allele frequency < 0.01 in western chimpanzees, > 0.9 in central chimpanzees, and fixed in bonobos.

Our study highlights the importance of terminological precision, such as whether allele frequencies are calculated using ancestral or derived alleles, and technical transparency, such as how ancestral alleles are inferred. Ambiguous definitions can lead to inconsistent interpretations and even propagation of errors in subsequent research. Likewise, the absence of publicly available implementations hinders reproducibility and independent validation. When methodological descriptions are unclear—for example, in the handling of ancestral alleles without explicit documentation—a well-documented implementation, such as a workflow built with Snakemake (Mölder et al. 2021), becomes an essential complement, clarifying the intended computational procedures. As interest grows in applying advanced computational methods, particularly machine learning and deep learning (Schrider and Kern 2018; Christin et al. 2019; Andermann 2020; Morimoto et al. 2021; Lürig et al. 2021; Borowiec et al. 2022; Callier 2022; Pichler and Hartig 2023; Korfmann et al. 2023; Yelman and Jay 2023; Huang et al. 2024; Mo et al. 2024; Wu 2024), to evolutionary biology, it is essential to ensure both rigorous theoretical formulations and accessible, reproducible implementations to uphold the reliability and integrity of scientific research (Huang 2024).

## Acknowledgements

We thank Florian Schmidt and Serhat Inceer for testing the SAI package. We also acknowledge the Life Science Compute Cluster at the University of Vienna for providing computing resources. This project has been funded by the Vienna Science and Technology Fund (WWTF) [10.47379/VRG20001] to M.K.

## Author contributions

X.H. designed the study. X.H. implemented SAI. X.H., S.C., J.H., and M.K. analyzed the data and wrote the manuscript.

## Data availability

The 1000 Genomes Project Phase 3 dataset can be downloaded from, https://ftp.ncbi.nih.gov/1000genomes/ftp/release/20130502/, last accessed April 19, 2025. The Lithuanian genomes can be downloaded from https://data.mendeley.com/datasets/d2xt5hdm5j/2, last accessed April 19, 2025. The Altai Neanderthal genome can be downloaded from http://cdna.eva.mpg.de/neandertal/Vindija/VCF/Altai/, last accessed April 19, 2025. The Denisovan genome can be downloaded from http://cdna.eva.mpg.de/neandertal/Vindija/VCF/Denisova/, last accessed April 19, 2025. BED files specifying variants that passed quality filters in the Altai Neanderthal genome can be downloaded from http://cdna.eva.mpg.de/neandertal/Vindija/FilterBed/Altai/, last accessed April 19, 2025. BED files specifying variants that passed quality filters in the Denisovan genome can be downloaded from http://cdna.eva.mpg.de/neandertal/Vindija/FilterBed/Denisova/, last accessed April 19, 2025. The bonobo and chimpanzee genomes can be downloaded from https://phaidra.univie.ac.at/pfsa/o_2066302/merged_segregating/Pan/Pan_wild_filtered/, last accessed April 19, 2025. The ancestral sequences based on the hg19 reference genome can be found in https://ftp.ensembl.org/pub/release-75/fasta/ancestral_alleles/homo_sapiens_ancestor_GRCh37_e71.tar.bz2, last accessed April 19, 2025.

The FASTA file for the reference genome hg38 can be found in https://hgdownload.soe.ucsc.edu/goldenPath/hg38/bigZips/hg38.fa.gz, last accessed April 19, 2025. The FASTA file for the reference genome rheMac10 can be found in https://hgdownload.soe.ucsc.edu/goldenPath/rheMac10/bigZips/rheMac10.fa.gz, last accessed April 19, 2025. The file for liftover chain from hg38 to rheMac10 can be found in https://hgdownload.soe.ucsc.edu/goldenPath/hg38/liftOver/hg38ToRheMac10.over.chain.gz, last accessed April 19, 2025. The source code of SAI can be found in https://github/xin-huang/sai, last accessed April 19, 2025. The Snakemake workflow for reproducing the analysis can be found in https://github.com/xin-huang/sai-analysis, last accessed April 19, 2025.

